# LEA 3: Factor models in population genetics and ecological genomics with R

**DOI:** 10.1101/2020.10.06.327783

**Authors:** Cléement Gain, Olivier François

## Abstract

A major objective of evolutionary biology is to understand the processes by which organisms have adapted to various environments, and to predict the response of organisms to new or future conditions. The availability of large genomic and environmental data sets provides an opportunity to address those questions, and the R package LEA has been introduced to facilitate population and ecological genomic analyses in this context. By using latent factor models, the program computes ancestry coefficients from population genetic data, and performs genotype-environment association analyses with correction for unobserved confounding variables. In this study, we present new functionalities of LEA, which include imputation of missing genotypes, fast algorithms for latent factor mixed models using multivariate predictors for genotype-environment association studies, population differentiation tests for admixed or continuous populations, and estimation of genetic offset based on climate models. The new functionalities are implemented in version 3.0 and higher releases of the package. Using simulated and real data sets, our study provides evaluations and examples of applications, outlining important practical considerations when analyzing ecological genomic data in R.

## 1 Introduction

Landscape and ecological genomics attempt to characterize geographic processes underlying the responses of organisms to their environments (Schoville et al., 2012; Sork et al., 2013; Manel & Holderegger, 2013; Savolainen, Lascoux, & Merilä, 2013). In these approaches, the recent availability of large genomic and environmental data sets have facilitated the identification of biotic and abiotic factors that influence neutral and adaptive genetic diversity patterns, offering opportunities for researchers to understand those patterns with statistical genetic approaches. Landscape databases include environmental variables such as climate and habitat descriptors which are proxies for geographically heterogeneous selection pressures (Fenderson, Kovach, & Llamas, 2020). Accounting for the confounding effects of demographic processes, local adaptation can be detected at a genomic level by identifying loci which allele frequency exhibits significant association with those environmental variables. Thus, ecological genomic studies could anticipate results from translocation experiments or exposure to future conditions by relying on analyses of population structure and genomic signatures of selection. Many methods and computer programs have been developed to this aim (Rellstab, Gugerli, Eckert, Hancock, & Holderegger, 2015; Hoban et al., 2016; Forester, Jones, Joost, Landguth, & Lasky, 2016), and the R package LEA – for *Landscape and Ecological Association* studies – is one of those programs (Frichot & François, 2015).

Methods in LEA are based on the statistical framework of latent factor models (Frichot, Schoville, Bouchard, & François, 2013; Frichot, Mathieu, Trouillon, Bouchard, & François, 2014). Latent factors are unobserved variables that represent data generated by processes linked to population history, population structure and technical or statistical artifacts. Latent representations of large data sets are usually computed as a reduced number of combinations of observed variables, a key step in statistics and in machine learning (Mardia, Kent, & Bibby, 1979; Murphy, 2012). Technically, the latent factor methods of LEA belong to the class of unsupervised machine learning approaches, and enable users to analyze population structure and detect genomic signatures of local adaptation without assumption on the biological processes that have generated the data. Several variants of latent factor models have been successfully applied in population genetic studies. Examples include estimates of ancestry coefficients with the Bayesian programs structure or tess, computations of eigenvectors in principal component analysis (PCA), uniform manifold approximation and projection for dimension reduction, and factor analysis for ancient DNA samples (Pritchard, Stephens, & Donnelly, 2000; Caye, Jay, Michel, & François, 2018; Patterson, Price, & Reich, 2006; Diaz-Papkovich, Anderson-Trocmé, & Gravel, 2019; François & Jay, 2020). In most applications of factor models, projections of individuals on factors reflect their levels of admixture from source populations. In genotype-environment association methods, latent factor regression models were introduced to separate variation explained by observed environmental variables from variation explained by unobserved variables (Frichot, Schoville, Bouchard, & François, 2013). In those regression models, latent factors represent unobserved confounders, and they have less direct interpretations than in ancestry estimation methods. Machine learning algorithms that train models to estimate latent factors are generally computationally efficient with minor loss of statistical accuracy compared to Bayesian Monte-Carlo methods. Thus, they allow their users to increase the volume of data analyzed compared to previous approaches (Caye, Jumentier, Lepeule, & François, 2019).

In this study, we present functionalities implemented in LEA version 3 to perform imputation of missing genotypes, improved estimation in latent factor mixed models (LFMM), genome scans for selection based on factor models, and prediction of genetic offset under future environments. Here, future environments should be interpreted as new environments for which we have actual or projected data. Several algorithms processing high-throughput sequencing data do not accept missing genotypes, or handle these values with naive approaches, such as imputation with mean values. Missing data are problematic in genome-wide regression analyses, such as association studies, that have decreased power when genotypes are removed (Marchini & Howie, 2010). Missing data are also problematic for PCA when it is used for describing structure in the data. See Dray & Josse (2015) for imputation strategies related to PCA. Imputation of missing data has been an intensive field of statistical research for decades (Van Buuren, 2018). In population genetics, several methods have been proposed to address this issue based on reference genomes. For example, the fastphase model imputes missing genotypes by using linkage disequilibrium and hidden Markov models (Scheet & Stephens, 2006). Reference genomes are however not available for all organisms, and alternative methods relying on unsupervised machine learning have been considered (Stekhoven & Bühlmann, 2012; Chi, Zhou, Chen, Del Vecchyo, & Lange, 2013). LEA 3 implements an imputation algorithm based on factors estimated in the snmf function with a nonnegative matrix factorization approach (Lee & Seung, 1999; Frichot, Mathieu, Trouillon, Bouchard, & François, 2014). With this approach, missing genotypes are replaced by predicted genotypes in a way that agrees with the inference of population structure.

The R package LEA also implements algorithms for LFMMs, which are statistical models used in genotype-environment association studies to identify genomic signatures of adaptation to the local environment (Frichot, Schoville, Bouchard, & François, 2013). LFMMs optimally separate neutral genetic variation – modelled in the latent factors – from adaptive genetic variation modelled in the effect sizes of environmental covariates. Adaptive loci are expected to be associated with non-null effect sizes, that are tested at each locus in the genomic data. Frichot et al. (2013) used a Bayesian approach and a Markov Chain Monte Carlo (MCMC) algorithm to adjust LFMMs to the data, and this approach was implemented in the lfmm function of LEA (Frichot & François, 2015). Based on least-squares optimization methods (Caye, Jumentier, Lepeule, & François, 2019), much faster algorithms are now implemented in the lfmm2 function of LEA 3.

In addition to genotype-environment association methods, LEA 3 implements genome scans for selection based on population differentiation – also called outlier tests. Those tests screen a large number of genomic variants across a genome to identify loci that have been affected by diversifying selection. To achieve this objective, they consider the upper tail of the empirical distribution for population differentiation statistics like fixation indices and related measures (Lotterhos & Whitlock, 2015; Duforet-Frebourg, Luu, Laval, Bazin, & Blum, 2016; François, Martins, Caye, & Schoville, 2016). In this context, admixed individuals and genetically continuous populations complicate the use of population differentiation tests. Extensions of population differentiation measures have been recently proposed for samples with admixed individuals (Martins, Caye, Luu, Blum, & François, 2016). LEA 3 implements the statistics introduced by Martins et al. (2016), and computes significance values by comparing the values of those statistics with the genomic background.

Understanding the vulnerability of species and populations to environmental threats is important for developing effective strategies to conserve them (Foden et al., 2019). For a target species, genomic data can be used to evaluate evolutionary potential for future adaptations within a fixed time horizon (Sork et al., 2010; Jay et al., 2012; Pauls, Nowak, Bálint, & Pfenninger, 2013; Aitken & Whitlock, 2013; Razgour et al., 2019). The genomic approaches exploit the correlation of allele frequencies with geographical variation of the environment to predict mismatches between current and future allele distributions under environmental change projections (Fitzpatrick & Keller, 2015; Rellstab et al., 2016; Capblancq, Fitzpatrick, Bay, Exposito-Alonso, & Keller, 2020). In most applications, a set of adaptive loci is first identified from the genomic background using a genotype-environment association method. Then this set of adaptive loci is used to evaluate genetic offset based on another statistical approach (Bay et al., 2018; Ingvarsson & Bernhardsson, 2020; Waldvogel et al., 2020). LEA 3 implements an approach assuming that effects of the environment are potentially weak but highly polygenic. Using effect sizes instead of significance values, the new approach leverages the entire set of genotyped loci to predict genomic variation under projected conditions. Several examples and simulation studies illustrate the functionalities of LEA 3 below.

## 2 New program functionalities

### Imputation of missing data

LEA 3 implements an imputation method based on allele frequencies and ancestry coefficients estimated from its snmf function (Frichot, Mathieu, Trouillon, Bouchard, & François, 2014). Assuming *n* diploid organisms genotyped at *L* loci, the snmf algorithm decomposes the *n* × *L* matrix of observed allele frequencies, *P*, in a product of two probabilistic matrices

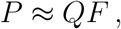

where the coefficients of *P* take their values in {0, 1/2, 1} for diploid organisms (Note: ploidy can be modified in snmf). The matrix *Q* is similar to the *Q*-matrix of STRUCTURE, representing ancestry coefficients for individuals originating from *K* source populations (Pritchard, Stephens, & Donnelly, 2000). The matrix *F* contains allele frequencies at each locus for each source population. While *P* may contain some missing values, the product matrix, *QF*, is always a complete probabilistic matrix. Thus imputation of missing genotypes can be achieved by replacing the missing values with random values sampled from the product matrix distribution. In the same spirit, an alternative option is to pick the most probable genotype based on the largest probabilities in a deterministic fashion. Both random and deterministic procedures were implemented in the impute function of LEA 3.

To evaluate the ability of the impute function to correctly reconstruct missing genotypes, we performed a simulation study of a two-population model with admixture. The simulation model assumes that a haploid population was genotyped at *L* diallelic loci (values 0 and 1) according to the *F*-model (Balding & Nichols, 1995; Pritchard, Stephens, & Donnelly, 2000). In the *F*-model, there is an ancestral gene pool containing alleles of unknown frequencies, *p*, having a uniform distribution over the *L* loci. At a given time point in the past, the ancestral population split in *K* sub-populations (here, *K* = 2), and the subpopulations diverged with genetic drift equal to *F*. Conditional on *p*, the allele frequency at a particular locus in any subpopulation follows a beta distribution of shape parameters *p*(1–*F*)*/F* and (1–*p*)(1–*F*)/*F*. The drift parameter, *F*, controls the fixation index, *F*_ST_, with larger values of *F* causing *F*_ST_’s to be larger. In simulations, the drift parameter was varied between 2% and 30%. In the model, a third population containing admixed genotypes was also sampled. The admixture rate *π*_admix_ was varied between 5% and 50%. For *n* = 300 individuals, the number of genotypes, *L*, was varied from 1,000 to 10,000. To create missing genotypes, we removed genotypes at random at a rate, *π*_missing_, between 10% and 90%. Ten runs of snmf were performed for each genotypic data set. The best run was retained on the basis of the cross-entropy criterion, and was used for imputation of the missing data. Then we checked whether the removed genotypes were correctly predicted and restored by the deterministic option of the program. The accuracy of reconstruction was measured as the percentage of correctly reconstructed genotypes, and with the root mean squared differences between ancestry estimates computed from the original and reconstructed genotypes. A total of 18,000 program runs for 1,800 data sets were performed. The program snmf was run with *K* = 2. The regularization parameter of snmf was increased to *α* = 100 (default value *α* = 10) to account for the relatively small number of loci in the simulations (Frichot, Mathieu, Trouillon, Bouchard, & François, 2014). Note that parameters used in simulations do not provide any guidelines for snmf in empirical studies. For example, it is always better to increase the number of runs. For their studies, users should refer to original method descriptions, choosing the regularization parameter (and *K*) based on the entropy criterion and cross-validation (Owen & Perry, 2009). Turning to real data, we also simulated missing genotypes in chromosome 5 for 162 European accessions of *Arabidopsis thaliana* (53,859 single nucleotide polymorphisms, SNPs) (Atwell et al., 2010). This was done by removing actual genotypes with a rate varying between 10% and 90%.

### Fast latent factor mixed models

LEA 3 (re)implements the LFMM ridge estimation algorithm presented by Caye, Jumentier, Lepeule, & François (2019). The lfmm2 function handles multivariate environmental data, and estimates locus-specific effect sizes and latent factors by using a least-squares method. The new implementation relies on core functions from the *base* package of R which warrants reproducibility of analyses on the long term. To compare lfmm2 with the MCMC algorithm programmed in lfmm, simulations were performed from the LFMM regression model. The model generates continuous values, and a deterministic link function was added in order to obtain haploid genotypes (negative values corresponded to allele 0 and positive values corresponded to allele 1). Following this model, one hundred genotypic matrices were created for *n* = 100 individuals genotyped at *L* = 2, 000 loci. True associations were simulated at 100 loci, with effect sizes between −10 and 10. A Gaussian environmental variable, *X*, was simulated for each individual, and population structure was modeled by *K* = 3 Gaussian latent factors, exhibiting various levels of collinearity with the environmental variable. More precisely, collinearity between *X* and the latent factors was measured by the coefficient of determination, which varied between 10% and 80% in the simulations. High levels of collinearity corresponded to strong confounding effects, and were expected to decrease the power of association tests (Frichot, Schoville, de Villemereuil, Gaggiotti, & François, 2015b). For the lfmm and lfmm2 tests, power and false discovery rate (FDR) were computed after a Benjamini-Hochberg procedure was applied with an expected FDR level of 5% (Benjamini & Hochberg, 1995). Power was computed as the proportion of true associations that are correctly discovered, and FDR was computed as proportion of false discoveries among the discoveries. In LFMM analyses, the number of factors was set equal to *K* = 3, corresponding to a recommended value obtained from the elbow in PCA scree-plots for the genetic data.

In a second series of experiments, we compared the relative performances of lfmm2 with genome scans based on redundancy analysis (RDA) (Mardia, Kent, & Bibby, 1979; Forester, Lasky, Wagner, & Urban, 2018). To this objective, we re-analyzed one hundred simulated data sets from Capblancq, Luu, Blum, & Bazin (2018). Each data set was obtained from forward simulations of biologically realistic scenarios with weak population structure, and contained *n* = 640 diploid individuals genotyped at *L* = 1, 000 SNPs. In these scenarios, three quantitative traits were controlled by 10 unlinked loci (QTLs). Ten environmental variables were also created. The first three environmental variables determined selective pressure on the traits, whereas the other ones had no effect on the phenotypes. Power and FDR were computed for lfmm2, RDA and partial RDA. Partial RDA was introduced to correct for confounding due to population structure and to provide a fair comparison of algorithms by conditioning RDA on latent factors estimated by lfmm2. To compare the results with (Capblancq, Luu, Blum, & Bazin, 2018), we used the parameters and the computer codes of their study, performing association analyses with principal components built on environmental variables and setting the expected FDR level to 10%.

### Identifying outlier loci with latent factor models

Statistical tests to identify loci associated with extreme values of population differentiation statistics were implemented in the snmf.pvalues function of LEA 3 (Martins, Caye, Luu, Blum, & François, 2016). For *K* ancestral populations, the population differentiation statistics were defined at each locus as follows

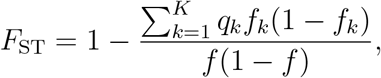

where *q*_*k*_, *f*_*k*_ and *f* were obtained from the *Q* and *F* estimates computed by the snmf function. The quantity *q*_*k*_ was equal to the average ancestry coefficient across individuals, *f*_*k*_ was equal to the allele frequency in population *k*, and we set 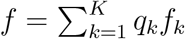. Based on those modified *F*_ST_ statistics, significance values were computed by using the Fisher distribution. Fisher tests are similar to chi-squared tests, which have been extensively studied both for snmf and for PCA (Martins, Caye, Luu, Blum, & François, 2016; Duforet-Frebourg, Luu, Laval, Bazin, & Blum, 2016; Chen, Lee, Zhu, Benyamin, & Robinson, 2016; Galinsky et al., 2016). With PCA, genome scans were performed by fitting a regression model for each SNP, for which the response variable is the SNP frequency and the explanatory variables are the *K* – 1 first PCs of the genetic data. We illustrated the results of outlier tests implemented in the snmf.pvalues function by re-analyzing SNP data for 49 accessions from Scandinavian lines of *Arabidopsis thaliana* (Atwell et al., 2010). Filtering loci for minor allele frequency greater than 5% resulted in 205,417 SNPs across the five chromosomes. We compared the results of the tests with tests based on PCA loadings and with LFMMs considering latitude as an explanatory variable which correlates with population structure (*K* = 2).

### Genetic offset statistics

LEA 3 allows computing predictive measures of genetic offset based on future environmental data. This section presents a brief outline of the theory underlying the computation of genetic offsets, sufficient for the interpretation of the program outputs. For a single population, the offset statistic measures the divergence between allele frequencies in current conditions and in a fictive population harboring frequencies corresponding to future (or new) conditions. For this population, the function considers two sets of environmental variables, **X**_current_ and **X**_future_, for current and projected individual environmental conditions. The matrix of current variables is first used to fit an LFMM, and the fitted model is then applied to the new data for the prediction of responses. Technically, two genetic matrices, **Y**_fit_ and **Y**_pred_ are constructed

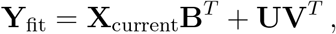

and

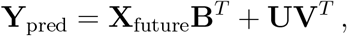

where **B**, **U**, and **V** are the effect size, factor and loading matrices adjusted by the lfmm2 algorithm from the current data. In matrix notation, **B**^*T*^ denotes the transpose of the matrix **B**. Then we considered *σ*_pred_ and *σ*_fit_ the largest singular values of the matrices **Y**_pred_ and **Y**_fit_, and *σ*_pred+fit_ the largest singular value of the concatenated matrix (**Y**_pred_, **Y**_fit_)^*T*^. The singular values were computed with the svd function of R, after all matrices were standardized. A *genetic offset*, *F*_offset_, is computed as follows

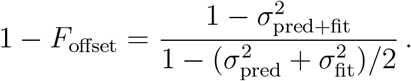

This definition of genetic offset is justified by the spectral analysis of population structure (Patterson, Price, & Reich, 2006). According to the genealogical interpretation of singular values (McVean, 2009; Bryc, Bryc, & Silverstein, 2013), the quantity *F*_offset_ is similar to a drift coefficient in a two-population model (Slatkin, 1991). The genetic offset measures the amount of genetic drift separating the population adapted to the current range of environmental variables to the fictive population adapted to the range of projected variables. In comparison to existing statistics (Fitzpatrick & Keller, 2015; Rellstab et al., 2016), the new measures do not select a particular subset of outlier loci but instead integrate over effects of environment at the genome scale, accounting for population structure. The statistics were implemented in the genetic.offset function of LEA 3.

To illustrate the function, we considered a subset of the 1,001 Genomes data for the plant species *Arabidopsis thaliana* (Alonso-Blanco et al., 2016). Two-hundred forty-one accessions from Southern, Central and Northern Sweden were extracted from the database. A matrix of SNP genotypes was obtained by considering variants with minor allele frequency greater than 5% and a density of variants around one SNP every 500 bp (334,946 SNPs). Because the genetic offset is a population statistic, we needed to cluster the individuals into groups. The individuals were clustered in eight groups after a preliminary analysis of population structure with snmf and on the basis of geographic proximity. To account for residual population structure in each of the eight groups, we chose *K* = 4 factors in the LFMM predictions (a conservative choice having minor impact on the results). Global climate and weather data corresponding to individual geographic coordinates were downloaded from the WorldClim database (http://worldclim.org). Eighteen bioclimatic variables, derived from the monthly temperature and rainfall values, were considered as representing the current environmental matrix. Projected environmental variables were obtained from three Representative Concentration Pathway (RCP) trajectories adopted by the IPCC fifth Assessment Report (2014). The pathways described different climate futures, corresponding to RCP 2.6, RCP 4.5, and RCP 8.5 (70 years) (IPCC, 2014).

## 3 Results

### Imputation of missing data

To evaluate the ability of the impute function to correctly reconstruct missing genotypes, we analyzed simulated genotypes from two populations with divergence and admixture. Over all data sets, the reconstruction accuracy for a missing genotype was around 76.5%, with a standard deviation of 14.1% (Figure 1). It was similar to the accuracy of reconstruction of non-missing genotypes by matrix factorization (mean around 77.5% with a standard deviation of 12.3%). The root mean squared differences between ancestry coefficients estimated from true and imputed genotypes were of the same order as the root mean squared differences between ancestry coefficients computed from distinct runs on the full data set (mean = 5.2% with a standard deviation of 3.1%). These results show that the reconstructed matrices were statistically similar to the original ones having no missing data. Significant association of accuracy with the proportion of missing data was observed (Multiple *R*-squared: 0.27, *F*-statistic: 660.3 on 1 and 1798 df, *P* lower than 2.2e-16). The results indicated that imputation was robust to removing a large fraction of genotypes, up to 30% (Figure 1A). Significant association was also observed between accuracy and the drift parameter (Multiple *R*-squared: 0.65, *F*-statistic: 3291 on 1 and 1798 df, *P* lower than 2.2e-16), indicating that imputation performed better when the population split occurred at older dates (Figure 1B). There were weak effects of the number of loci considered and admixture rate, leading to non-significant *p*-values. We eventually tested the effect of interaction between the simulation parameters, and found that accuracy was best explained by the following combination of simulation parameters (Multiple R-squared: 0.92, F-statistic: 5125 on 4 and 1795 df): 0.747 + 0.221 *F* - 0.010 *π*_missing_ - 0.106 *F* × *π*_missing_ - 0.073 *F* × *π*_admix_. Accuracy increased with larger values of *F* but not so much when the rate of missing values or the admixture rate increased at the same time. Next, we considered missing genotypes in European accessions of *Arabidopsis thaliana* by removing a proportion *π*_missing_ of actual genotypes. Using a model with *K* = 6 clusters, the accuracy of reconstructed genotypes varied approximately as 0.784 - 0.009 *π*_missing_ − 0.050 *π*^3^_missing_ (Multiple *R*-squared: 0.99). For 10% missing values, the total number of correct genotypes in the imputed data was around 98% of the *Arabidopsis* data, which had no missing genotypes. For 80% missing values, the total number of correct genotypes in the imputed data was around 80% of the original data.

**Figure 1.**
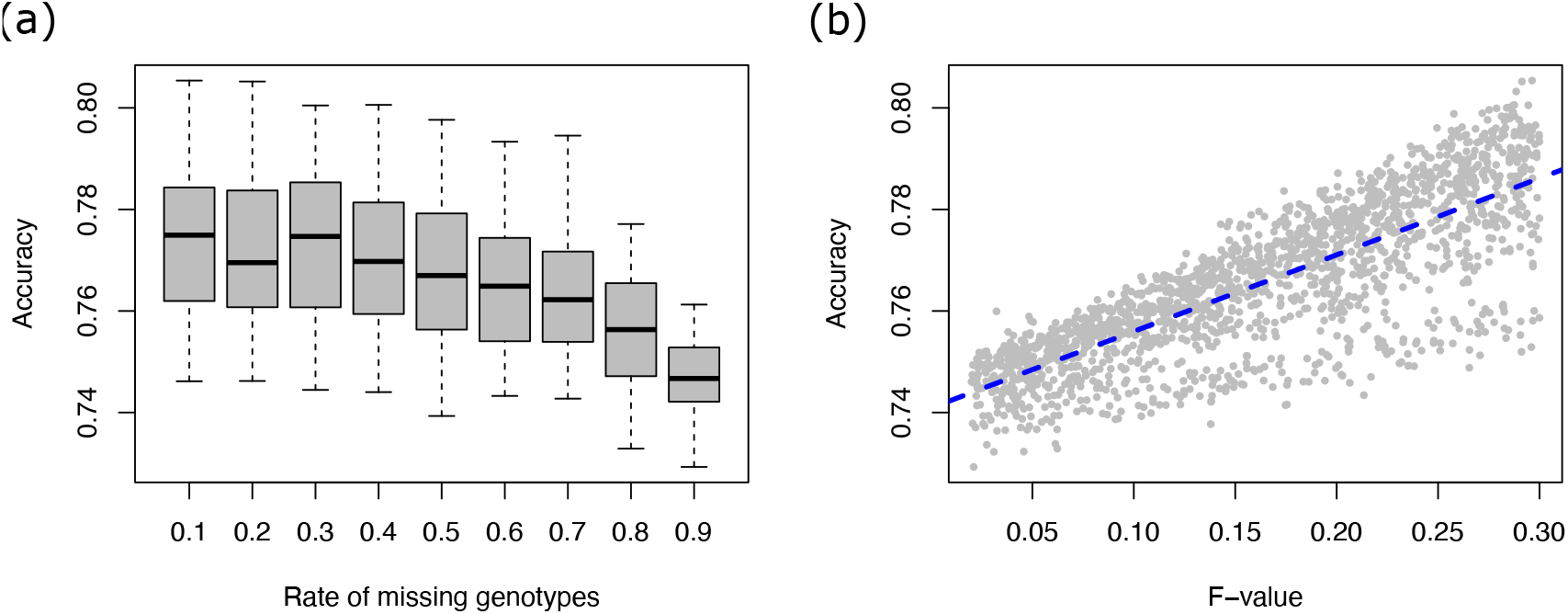
Imputation of missing data. Simulations from a two-population *F* - model with admixture. **a.** Accuracy of reconstructed genotypes as a function of the rate of missing genotypes. **b.** Accuracy of reconstructed genotypes as a function of the drift parameter *F*.

### Latent factor mixed models

To compare the statistical performances of the lfmm and lfmm2 estimation algorithms, 100 genotypic matrices were simulated, based on the common generative model for both approaches (Figure 2). The average FDR was around its expected value of 5% for lfmm (mean value 6.4%, Figure 2A). The average FDR was significantly lower for lfmm2 (mean value 3%, *t*-test = 6.88, *P* = 1.2e-10), showing that the least-squares algorithm was more accurate than the MCMC algorithm when simulations were performed under the generative model. For those simulations, the two versions had equivalent power to reject loci with null effect sizes (Figure 2B). Power was generally high (greater than 90%) when collinearity between population structure and the environmental variable was moderate, corresponding to values of the coefficient of determination lower than 60%. A sharp decrease in power was observed when the coefficient of determination reached values higher than 60%.

**Figure 2.**
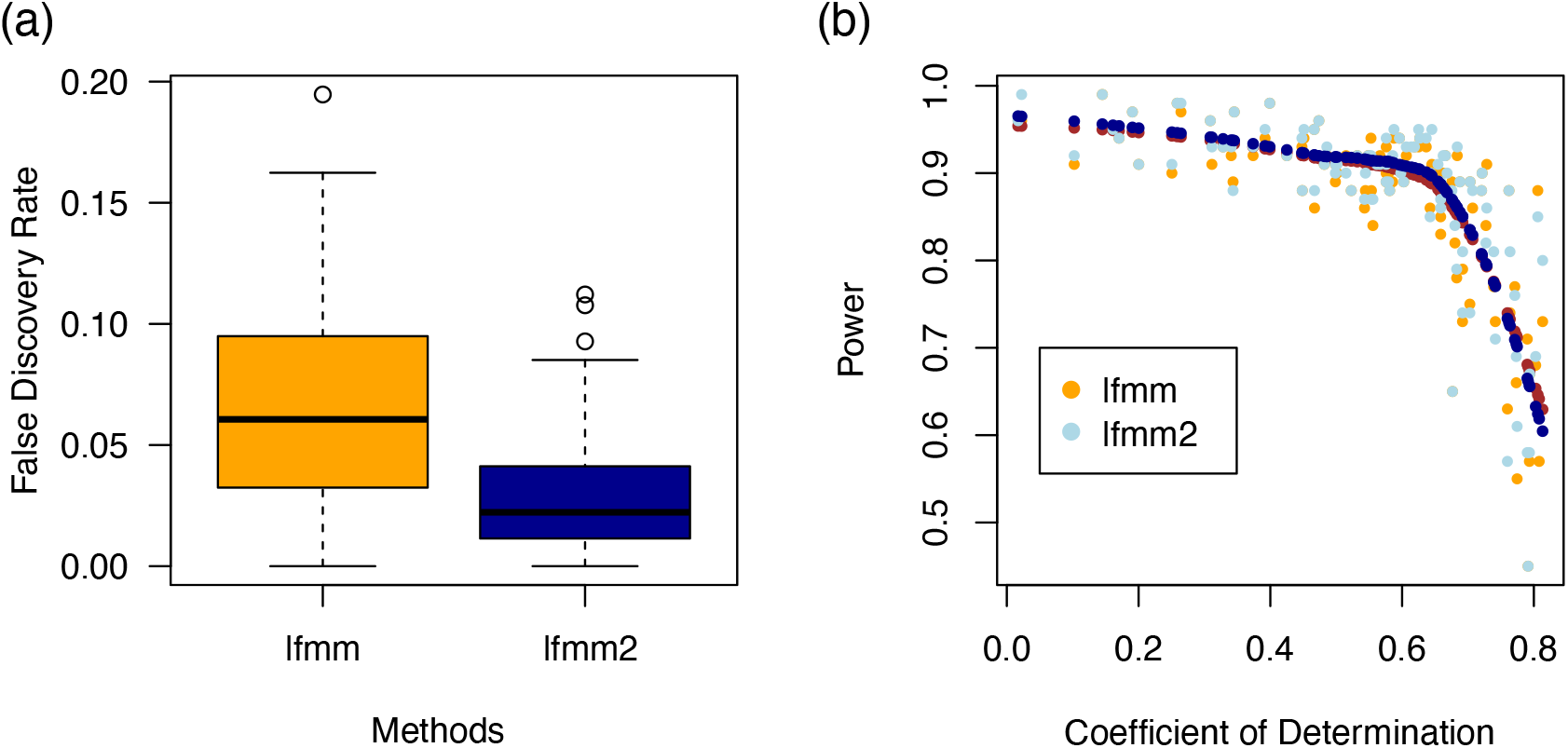
Comparisons of lfmm (MCMC) with lfmm2 (least squares). **a.** False discovery rates observed for an expected FDR of 5%. **b.** Power to reject null effect sizes. The coefficient of determination is the proportion of environmental variation explained by the hidden factors in the simulated data. The dark blue and (hidden) brown points correspond to local regressions of the power values.

Next, we compared the relative performances of lfmm2 with approaches based on RDA and partial RDA, resuming simulations conducted in (Capblancq, Luu, Blum, & Bazin, 2018) (Figure 3). Power and FDR obtained with RDA and partial RDA were highly similar, in agreement with weak population structure observed in the simulations (Capblancq, Luu, Blum, & Bazin, 2018). Power was close to 100% for all 10 SNPs in QTL2 with all methods. FDRs were around their expected values for all methods (10%). Overall, lfmm2 had increased power compared to RDA approaches for SNPs in QTL1 (mean value for lfmm2 = 90%, mean value for RDA = 75%, *t*-test = 8.01, *P* = 1.602e-13) and QTL3 (mean value for lfmm2: 50%, mean value for RDA: 20%, *t*-test = 11.45, *P* lower than 2.2e-16), contrasting with results for lfmm reported in the previous study.

**Figure 3.**
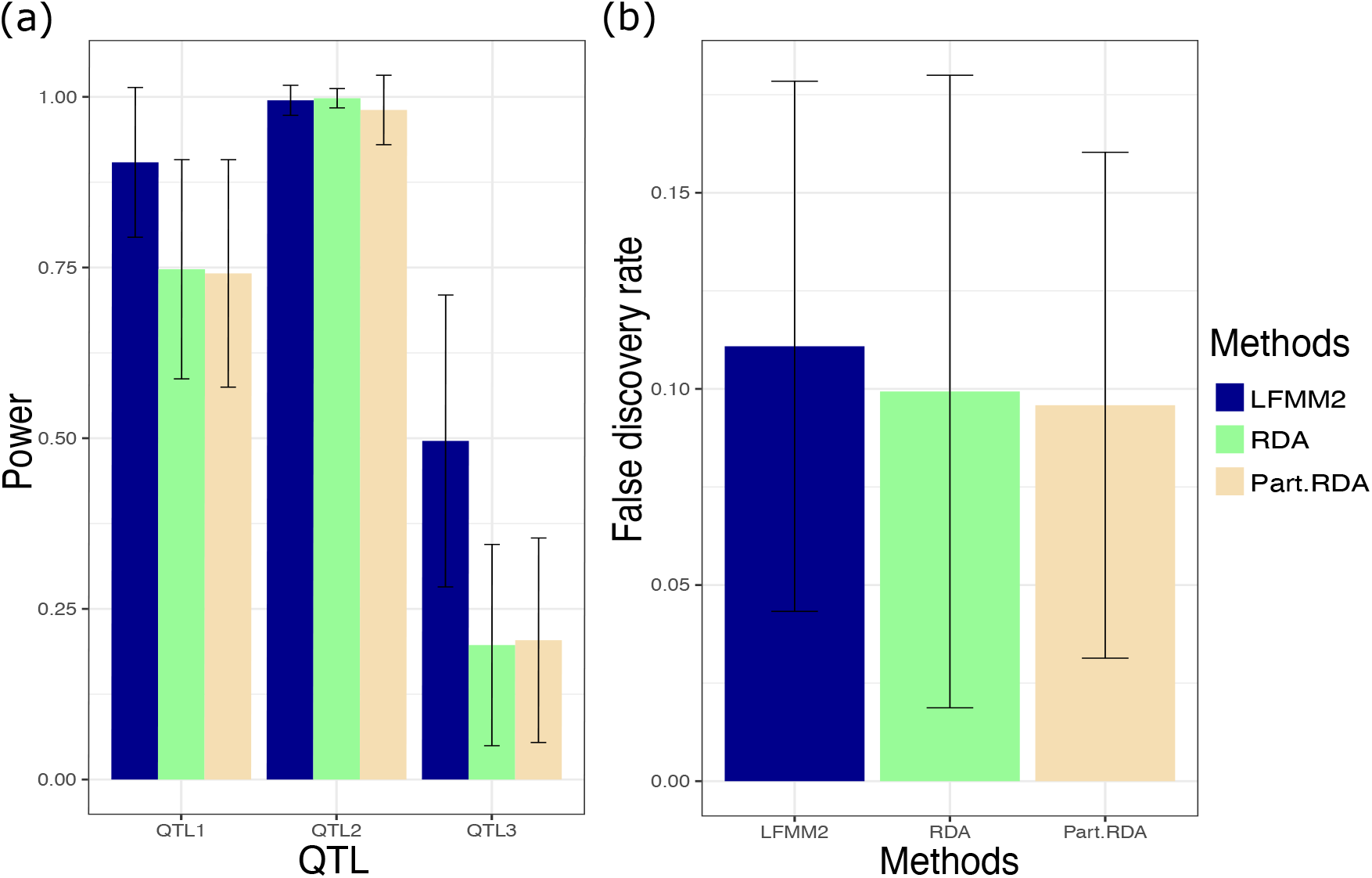
Comparisons of lfmm2 with RDA and partial RDA. Simulations resumed from (Capblancq et al. 2018). **a**. Power to detect loci under selection, **b.** False discovery rate for a controlled FDR level of 10%. Partial RDA (Part. RDA) includes corrections based on LFMM latent factors.

### Identifying outlier loci with latent factor models

We illustrated outlier tests based on population differentiation statistics introduced in Martins, Caye, Luu, Blum, & François (2016) by re-analyzing SNP data for 49 Scandinavian accessions *Arabidopsis thaliana* with snmf.pvalues, with PCA loadings and with lfmm2. The samples were divided in two geographic groups located in southern (37 individuals) and in northern Sweden (12 individuals). Latitude was used as an environmental variable in lfmm2. Although the histograms of *P*-values exhibited similar shapes for all methods (Figure S1), differences were observed in the tails of the test statistics. The *P*-values were lower in snmf than in lfmm2 and PCA (Figure 4). The Pearson correlation between snmf and PCA significance values was high, around 95%, and greater than the correlation between PCA and lfmm2 values (91%). These results can be explained by a high correlation between ancestry coefficients and PC1 scores (Figure S2). Despite the differences in the tails of distributions, genome scans based on snmf, PCA and lfmm2 hit the same genomic regions in chromosomes 1 and 5 of the plant, and the ranking of association levels were comparable in the three approaches. After Bonferroni correction, there were 1,013 significant hits at the 5% nominal level for snmf, 485 for PCA, and 459 for lfmm2, corresponding to around 50 genomic regions (Figure 4). The top hits for snmf, with significance values above 50, did not reach the highest levels in the PCA and LFMM tests. Those 129 hits corresponded to SNPs with allelic frequencies close to fixation either in the southern or the northern group (Figure S3). Focusing on a region of chromosome 5 around 113 Mb which was enriched in top hits, we estimated heterozygozity along the chromosome (Figure S4). This analysis provided evidence that a few loci exhibit signatures of selective sweep, more often in the northern group than in the southern group.

**Figure 4.**
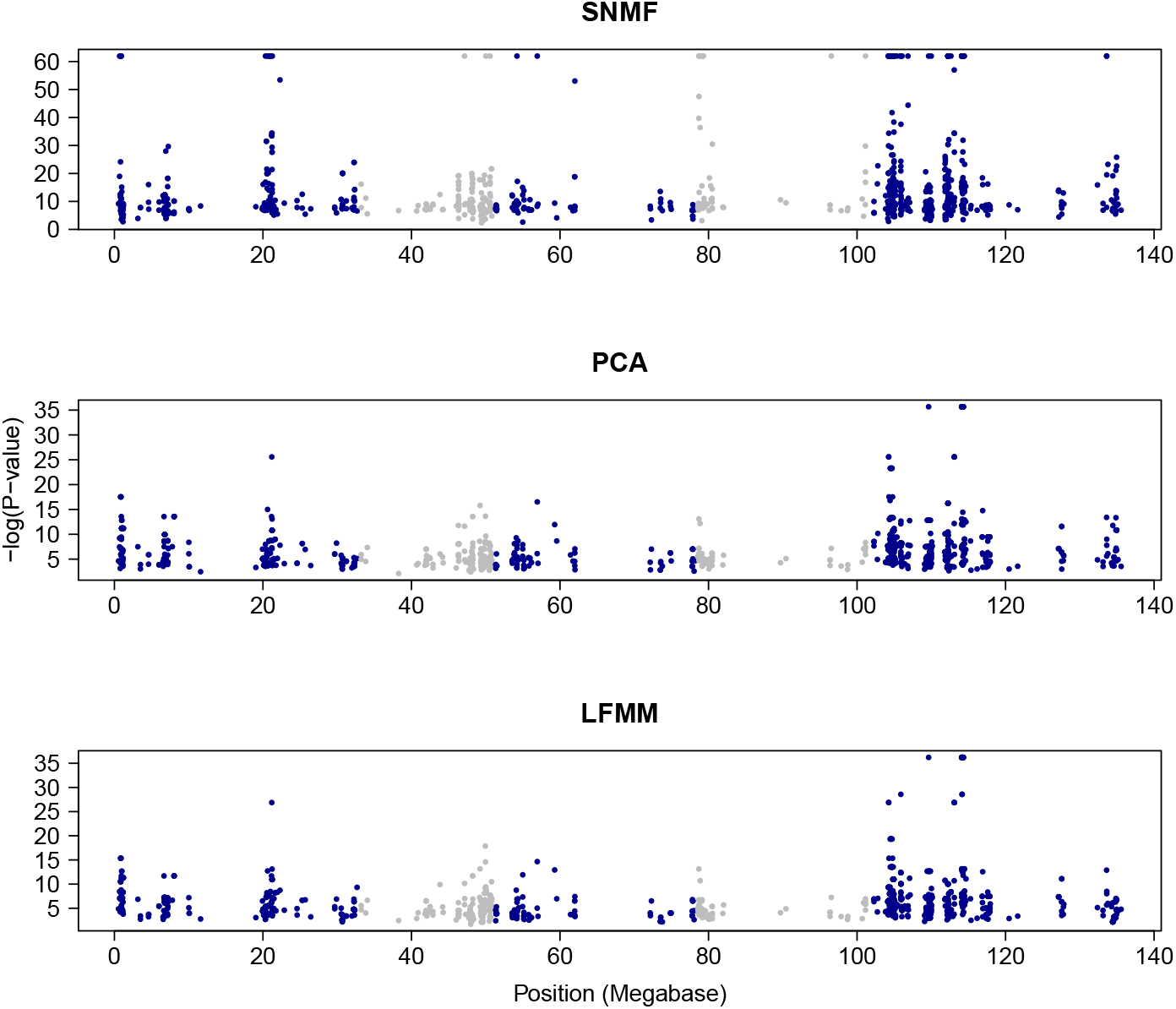
Genome scans for selection. SNP genotypes of forty-nine accessions of *Arabidopsis thaliana* from Northern and Southern Sweden analyzed with snmf, PCA, and lfmm2 with latitude used as an explanatory variable. Only loci with −log *P* greater than the Bonferroni threshold in one method (= 6.61) are shown. See Figure S4 for a focus on the region around 113Mb.

### Genetic offset statistics

Leveraging multivariate environmental data analysis with lfmm2, we illustrated the computation of genetic offset statistics for Scandinavian populations of *A. thaliana* (241 individuals) obtained from the prediction of eighteen bioclimatic variables based on three RCP trajectories (70 years). In LEA 3, genetic offset has the same interpretation as *F*_ST_, measuring the percentage of inbreeding in the population formed by the union of the current and projected samples. Under RCP 2.6, the genetic offset statistics ranged between 0% and 56%, with a mean value around 29% (Figure 5). For RCP 4.5 and 8.5, the maximum ranges stretched to 74% and 79% with mean values around 51% and 60% respectively (Figure 5). In RCP 2.6, the most exposed populations were at latitude around 50°N in Southern Scandinavia. Under RCP 4.5 and RCP 8.5, the most exposed populations were in the north at latitude around 62°-64°N. The risks were poorly explained by latitude alone (*R*^2^ less than 10%), but they were consistent with the principal components of projected climatic variables under the considered scenarios (PCs shown in Figures S5-S6).

**Figure 5.**
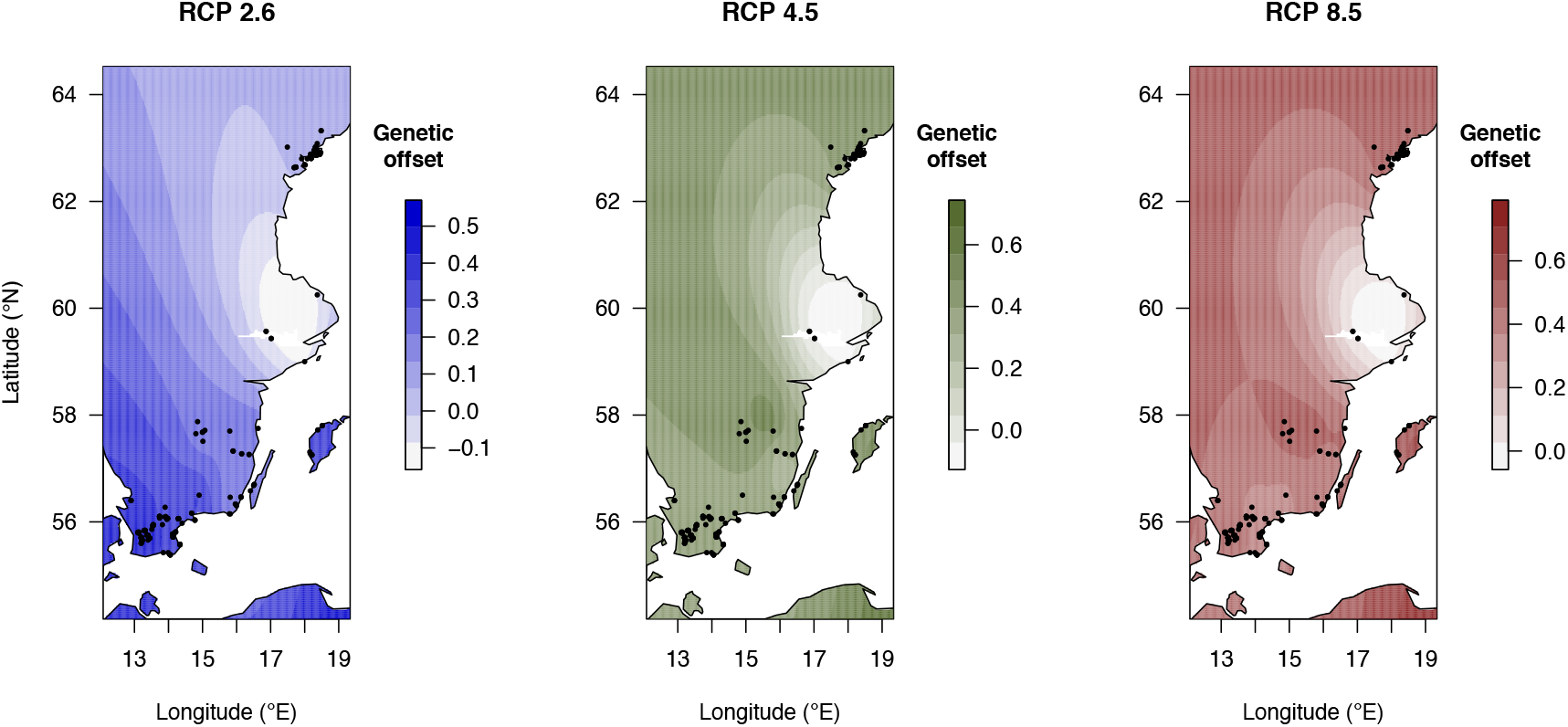
Genetic offset for Scandinavian populations of *Arabidopsis thaliana*. Genetic offset computed from projections of 18 bioclimatic variables according to three climate models RCP 2.6, 4.5 and RCP 8.5 (70 years). The genetic offset statistics were computed for 8 population samples. The values were interpolated by using the kriging algorithm implemented in fields 10.2 at the 241 sampling sites represented as black dots (Nychka et al. 2017).

## 4 Discussion

Modern molecular ecology studies make extensive use of unsupervised statistical methods for their analysis of large population genomic data (Hendricks et al., 2018; Paradis et al., 2017). The R program LEA provides an integrated suite of functions for running such analyses, including model-free inference of population structure and genome scans for signatures of local adaptation. This computer note presented functionalities implemented in version 3.1 of the computer program, and surveyed some applications of factorial methods in molecular ecology and ecological genomics. Since the first release of the package, new factor methods have been developed and integrated in the development version of the package. These new methods encompass genome scans based on population differentiation statistics and genotype-environment association studies, which have been evaluated in previous studies (Martins, Caye, Luu, Blum, & François, 2016; Caye, Jumentier, Lepeule, & François, 2019). The inclusion of those methods in LEA will facilitate their applications by R users, avoiding unnecessary coding or data formatting. Additional functionalities, including imputation of missing data and predictive values of genetic offset increase the range of applications of the package. A drawback of latent representations built in machine learning algorithms is to favor prediction to the detriment of interpretation (Lipton, 2018). With latent factor models, interpretation of results in terms of standard population genetic concepts, such as ancestry coefficients, population differentiation and genetic drift is preserved, as well as their appropriateness for predictive objectives.

Concerning missing data, the method implemented in LEA 3 is a novel approach, based on matrix factorization and on ancestry coefficients estimated by snmf (Frichot, Mathieu, Trouillon, Bouchard, & François, 2014). The matrix factorization technique has been frequently employed in machine learning applications where it has been successful in collective filtering and recommender systems (Koren, Bell, & Volinsky, 2009). The approach is conceptually similar to the imputation methods used by Bayesian programs like structure (Pritchard, Stephens, & Donnelly, 2000). Imputation based on snmf factors is consistent with respect to the allele frequencies in the ‘ancestral’ populations. Thus the method is appropriate for genotype-environment association studies and other applications using ancestral allele frequency information. An advantage of matrix factorization approaches is to be efficient without reference genomes. For example, imputation based on matrix factorization decreased the proportion of missing data from 10% to 2% in *Arabidopsis thaliana* simulations. While the limits on imputation are generally poorly understood and specific to each data set, we encourage users to remove a small proportion of non-missing genotypes (for example, 1%) and check the validity of their imputation results in preliminary analytical stages of their empirical studies.

Regarding genotype-environment association studies, least squares estimates for latent factor regression models were implemented in the lfmm2 function (Caye, Jumentier, Lepeule, & François, 2019). The function lfmm2 extends lfmm by allowing multivariate environmental data, which is particularly useful for predictive applications. The main improvements over lfmm are substantial savings of computer resources thanks to an exact algorithm rather than a Monte Carlo method. Although the lfmm2 and lfmm methods assumes slightly different sparsity conditions on effect size estimates, we found that they were equivalent in terms of sensitivity, reaching high levels of true positive rates when the coefficient measuring collinearity between environmental variables and latent factors was below 60%. An abrupt decrease in power was however observed for coefficients greater than 60%. These results suggests that users should attempt to evaluate the determination coefficient a priori from the first principal components of the genomic data or a posteriori from factors estimated by the methods. After reanalyzing simulated data sets from (Capblancq, Luu, Blum, & Bazin, 2018), we found that the multivariate version of lfmm2 compared favorably with genome scans based on redundancy analysis. The explanation for improvement over the results reported by Capblancq, Luu, Blum, & Bazin (2018) is that our analysis included more environmental information than did the previous study, exemplifying the benefit of using multivariate environmental data in LFMMs.

Population differentiation tests are often used in assessing genomic signatures of local adaptation. For *Arabidopsis thaliana*, population differentiation tests based on snmf, PCA and association with latitude highlighted similar genomic regions. Because those regression methods exhibit almost maximal collinearity between population structure and the explanatory variable, the interpretation of genome scans may be difficult, and it is complicated by the demographic history of Scandinavian *A. thaliana* populations (Huber, Nordborg, Hermisson, & Hellmann, 2014; Lee et al., 2017). A large fraction of the SNPs discovered by the genome scan methods might have resulted from neutral demographic processes or from linked selection. Choosing environmental variables not co-varying with population structure could alleviate the problem in genotype-environment association methods (François, Martins, Caye, & Schoville, 2016). With dense genomic data, we also suggest that population differentiation tests should be coupled with methods to detect selective sweeps (Vatsiou, Bazin, & Gaggiotti, 2016). For example, when we focused on a region enriched in snmf top hits, the analysis of heterozygozity along the fifth chromosome supported that some hits exhibited convincing signatures of selection, in particular in the northern group.

Following recent approaches in predictive ecological genomics (Bay et al., 2018; Fitzpatrick & Keller, 2015), LEA 3 can compute genetic offset statistics based on models of association between environmental variables and allele frequencies. The new statistic accounts for potentially small effects of the environment that are spread across the genome and the offset computation assumes that changes in population structure will occur only through modification of adaptive allele. In addition, genetic offset has an interpretation similar to a population differentiation statistic, measuring drift between allele frequencies in populations under current and future conditions. According to (Skoglund, Sjödin, Skoglund, Lascoux, & Jakobsson, 2014; Frachon et al., 2017), the values of this drift coefficient can be converted in generation time using the equation, *t* = 4*N*_e_*F*_offset_/(1 – *F*_offset_), where *N*_e_ is the effective population size. For Scandinavian populations of *A. thaliana*, relatively large values of *N*_e_ have been reported, around *N*_e_ ≈ 1, 000 (Lundemo, Falahati-Anbaran, & Stenøien, 2009). For these values, an offset statistic around 30-70% indicates that the evolutionary time required for adaptation to predicted climates will be around four thousands generations. While estimates may be biased upward by having removed rare variants from the analysis, a robust result is that genetic offset was higher in northern populations under the most extreme RCP scenario.

Computer programs evolve and upgrading LEA to include recent developments of latent factors in ecological genomics was necessary. The functions implemented in LEA 3 are built on classes of objects that are fully consistent with the previous versions of the program, so that users will adapt easily to the new version. Examples included in the program documentation and in a tutorial are available from www.bioconductor.org for new users to learn the program quickly.

## Supporting information

Sup. Mat.

## Acknowledgments

The authors are grateful to the organizers of the special issue for their invitation to submit a manuscript. They thank the reviewers for useful and constructive comments that improved the presentation of the manuscript. This work received support from the “Predictive Ecological Genomics” project (MIAI@Grenoble-Alpes, ANR-19-P3IA-0003).

## Data availability

The R package LEA 3.0 and higher versions are available from Bioconductor: www.bioconductor.org/packages/release/bioc/html/LEA.html. Development versions are available from www.bioconductor.org/packages/devel/bioc/html/LEA.html, and github.com/bcm-uga/LEA. The computer scripts and the data used in the examples or in the simulation studies of this manuscript are available from a GitHub repository github.com/bcm-uga/LEA3_simulation_script under GNU General Public License v3.0.

## Author contributions

OF designed research, CG and OF performed research, analyzed data and wrote the paper.

