## Supplementary material for "LEA 3: Factor models in population genetics and ecological genomics with R": Sup. Mat.

### Supplementary information for "LEA 3: Factor models in population genetics and ecological genomics with R"

Clément Gain Olivier François

**Authors' affiliations**:

Université Grenoble-Alpes, Centre National de la Recherche Scientifique, Grenoble INP, TIMC-IMAG CNRS UMR 5525, 38000 Grenoble, France.

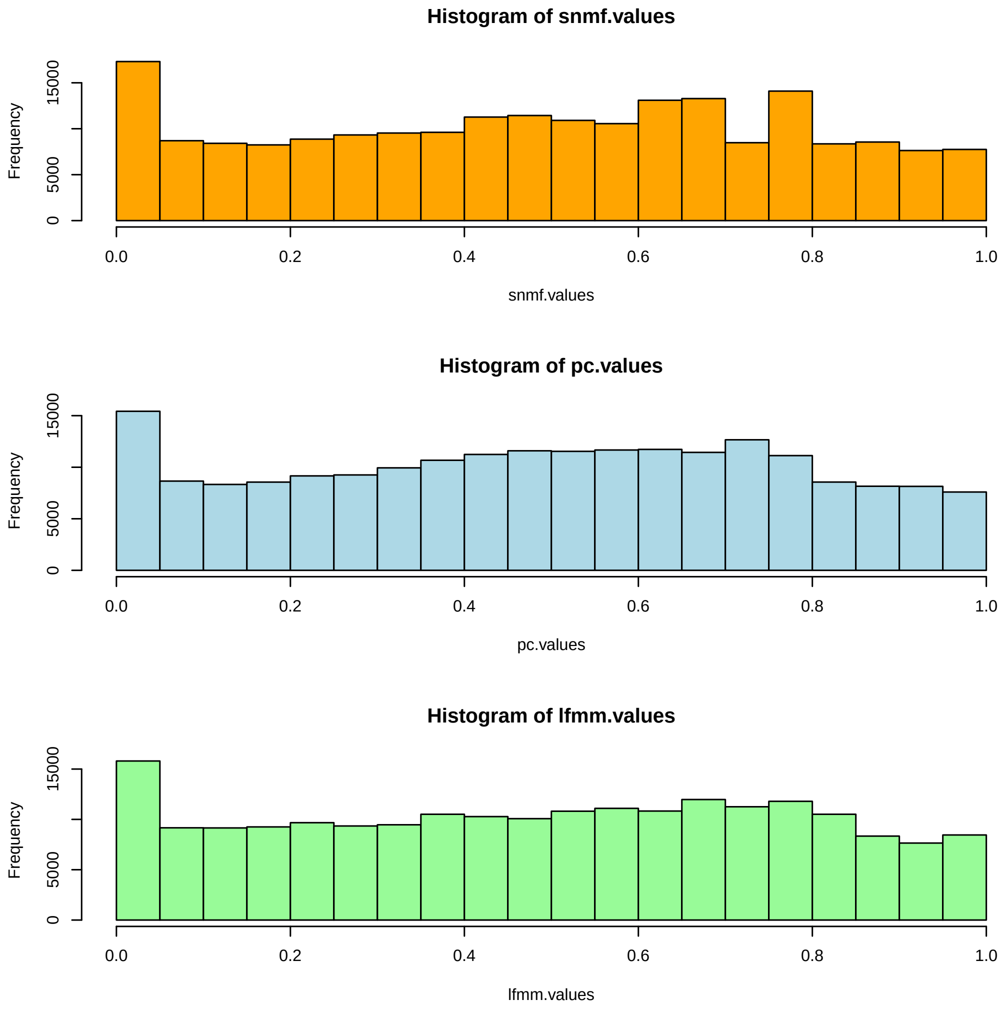


**Figure S1. Distribution of significance values for Scandivanian accessions of *A. thaliana*.** Histograms *P*-values for genome scans based on `snmf`, PCA, and `lfmm2` used with individual latitude information.

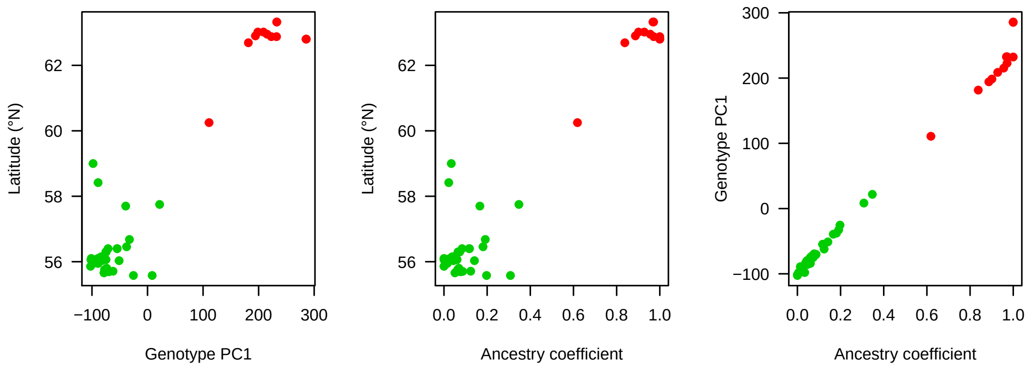


**Figure S2. Correlations between predictors in genome scans for Scandivanian accessions of *A. thaliana*.** Latitude (°N) for 49 plant accessions, PC1: first axis of a PCA performed on *A. thaliana* genotypes, Ancestry coefficients: admixture coefficients for the northern group estimated by `snmf` with $K = 2$. The colors show the membership of individuals to the northern (red) and southern groups (green).

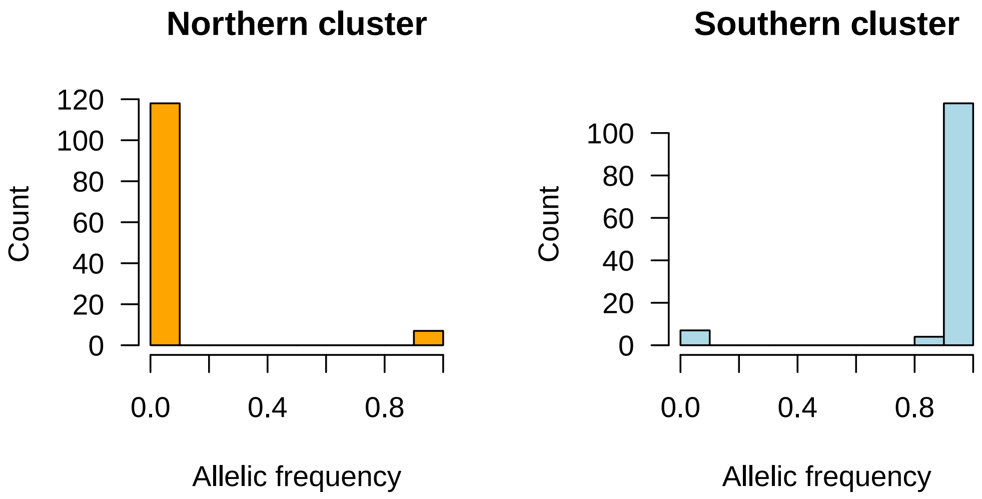


**Figure S3. Histograms of allelic frequencies for `snmf` top hits in a genome scan for Scandivanian accessions of *A. thaliana*.** Allelic frequencies were computed within clusters inferred by `snmf` with $K = 2$. The histograms show that the top SNPs are close to fixation in both clusters, and thus exhibit long range linkage disequilibrium.

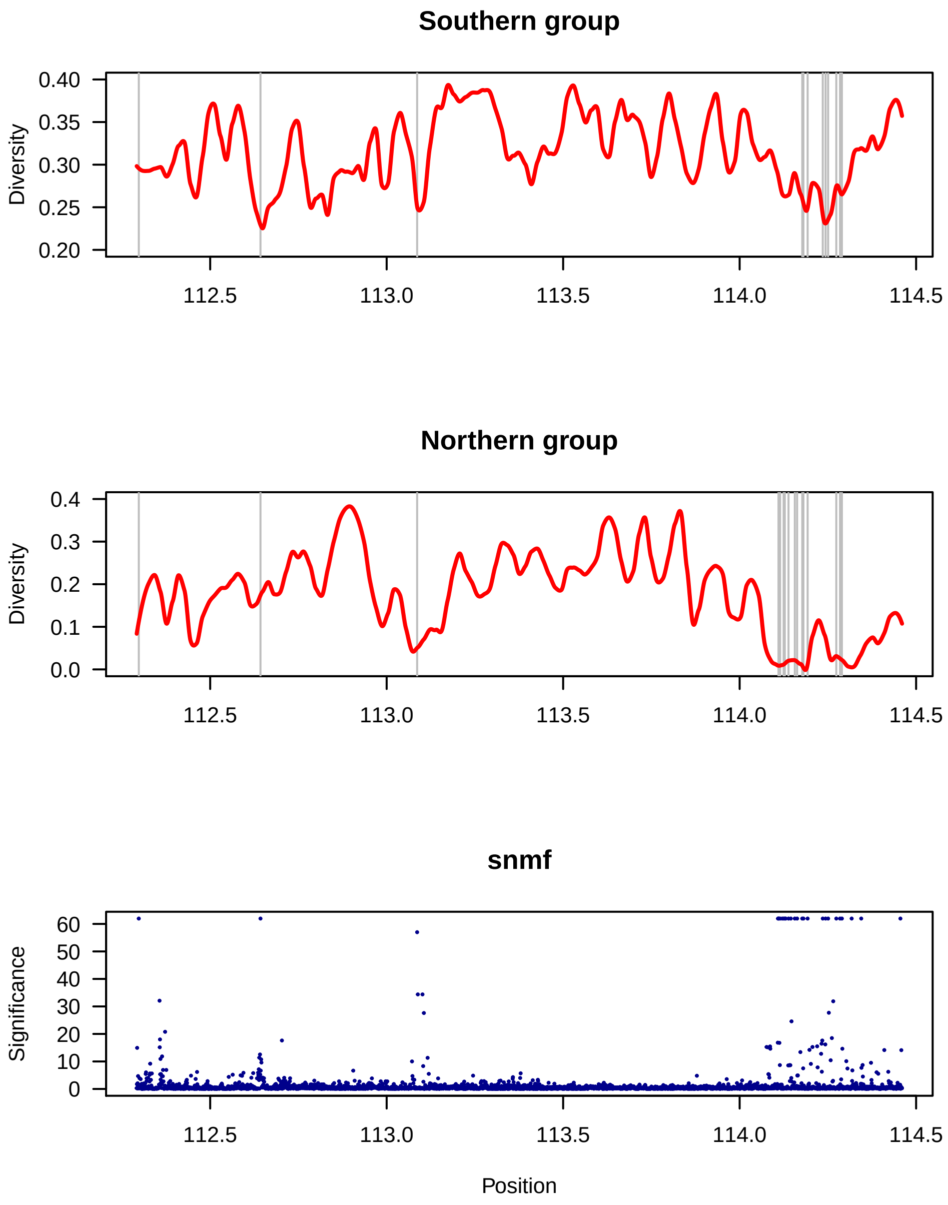


**Figure S4. Selective sweeps in the southern and in northern groups of *A. thaliana*.** Heterozygozity estimated in a region of the fifth chromosome centered at 113.5Mb for the southern ($n = 37$) and northern ($n = 12$) populations. The bottom row shows the significance values for `snmf` and grey bars correspond to the top hits for this method. Heterozygozity for a SNP with derived allele frequency $f$ was computed as $2f(1 - f)$ and adjusted to chromosomal position using a local regression method. The most remarkable signature of a selective sweep was around 114.2Mb in the northern group.

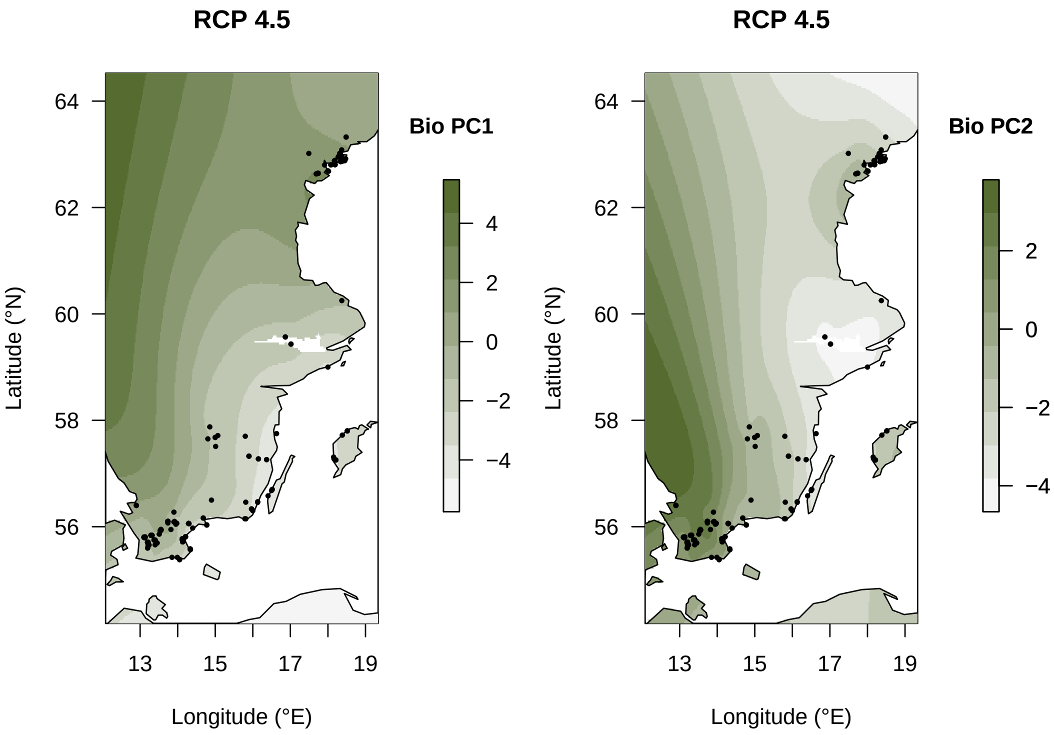


**Figure S5. Principal components of bioclimatic variables under RCP 4.5.** PC1 and PC2 of 18 bioclimatic variables predicted under climate model RCP 4.5 (70 years) interpolated at 241 sampling sites of Scandinavian *A. thaliana*.

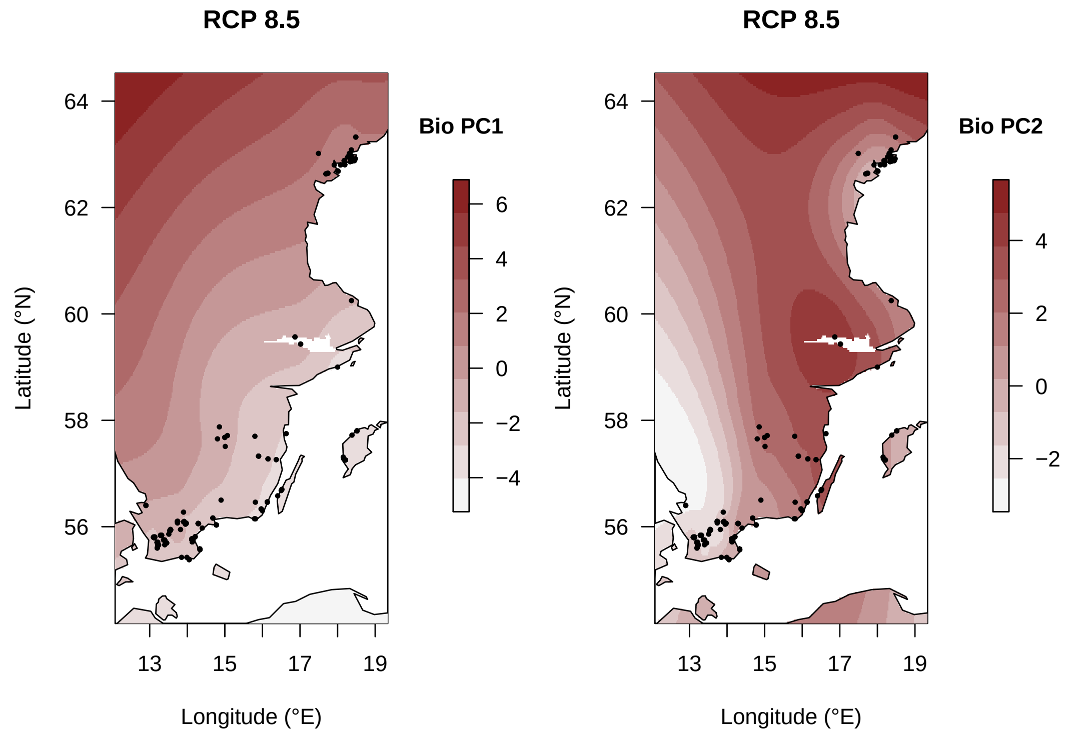


**Figure S6. Principal components of bioclimatic variables under RCP 8.5.** PC1 and PC2 of 18 bioclimatic variables predicted under climate model RCP 8.5 (70 years) interpolated at 241 sampling sites of Scandinavian *A. thaliana*.
